# Distinct neuronal subtypes of the lateral habenula differentially target ventral tegmental area dopamine neurons

**DOI:** 10.1101/743401

**Authors:** Michael L. Wallace, Kee Wui Huang, Daniel Hochbaum, Minsuk Hyun, Gianna Radeljic, Bernardo L. Sabatini

**Affiliations:** Howard Hughes Medical Institute, Department of Neurobiology, Harvard Medical School, Boston, MA 02115, USA.

## Abstract

The lateral habenula (LHb) is an epithalamic brain structure critical for processing and adapting to negative action outcomes. However, despite the importance of LHb to behavior and the clear anatomical and molecular diversity of LHb neurons, the neuron types of the habenula remain unknown. Here we use high-throughput single-cell transcriptional profiling, monosynaptic retrograde tracing, and multiplexed FISH to characterize the cells of the mouse habenula. We find 5 subtypes of neurons in the medial habenula (MHb) that are organized into anatomical subregions. In the LHb we describe 4 neuronal subtypes and show that they differentially target dopaminergic and GABAergic cells in the ventral tegmental area (VTA). These data provide a valuable resource for future study of habenular function and dysfunction and demonstrate neuronal subtype specificity in the LHb-VTA circuit.

## INTRODUCTION

The habenula is an epithalamic structure divided into medial (MHb) and lateral (LHb) subregions that receives diverse input from the basal ganglia, frontal cortex, basal forebrain, hypothalamus and other regions involved in processing both sensory information and internal state (Herkenham and Nauta, 1977; Yetnikoff et al., 2015). Two main targets of the LHb output are major monoaminergic structures in the brain, the ventral tegmental area (VTA) and raphe nuclei (dorsal (DRN) and medial (MRN)), whereas the MHb targets the interpeduncular nucleus (Herkenham and Nauta, 1979). Due to its dual effects on dopamine and serotonin producing neurons, LHb has been proposed to contribute to the neurobiological underpinnings of depression and addiction (Li et al., 2011, 2013; Maroteaux and Mameli, 2012; Meye et al., 2016). Furthermore, the LHb has been implicated in a wide range of functions and behaviors including reward prediction error, aversion, cognition, and adaptive decision making (Hikosaka, 2010; Matsumoto and Hikosaka, 2007; Mizumori and Baker, 2017; Proulx et al., 2014; Tian and Uchida, 2015; Wang et al., 2017).

The effects of LHb on each downstream structure and its contribution to different behaviors are likely carried out by distinct populations of neurons. In addition, the LHb may, at the macroscopic level, consist of distinct sub-domains with differential contributions to limbic and motor functions (Zahm and Root, 2017). Further, many studies suggest that habenular neurons show differences in gene expression across hemispheres (Concha and Wilson, 2001; Pandey et al., 2018), projection targets (Quina et al., 2014, 2015), and anatomical location (Gonçalves et al., 2012). Nevertheless, the systematic relationship between molecular profiles, projections patterns, and anatomical organization of neurons in the LHb is unknown.

Here we provide a comprehensive description of the neuronal subtypes in the LHb based on single-cell transcriptional profiling, multiplexed fluorescent *in situ* hybridization (FISH), and cell type specific monosynaptic retrograde tracing. Furthermore, as the MHb was included in our dissections, we also provide a molecular description of this nucleus. We find that the MHb has five, and LHb has four, transcriptionally-defined neuronal subtypes. Interestingly, the HbX subtype, which lies at the border between the MHb and LHb, is more transcriptionally similar to other LHb subtypes than MHb subtypes. We find that the four LHb neuronal subtypes are distinct and monosynaptic retrograde tracing revealed that they differentially target the dopaminergic and GABAergic neurons of the VTA. Furthermore, we find the LHb is organized into subregions defined by these transcriptionally discriminable neuronal subtypes. Together we identify previously unknown neuronal heterogeneity in the habenula and reveal that different neuronal classes have target biases in the VTA.

## RESULTS

### Cell type composition of the habenula by transcriptomic profiling

To examine cellular heterogeneity in the habenula, we performed high-throughput single-cell transcriptional profiling (“InDrop”) (Klein et al., 2015). Cell suspensions from the habenula were generated from acute, microdissected brain slices from adult mice (Figure 1A), producing 25,289 single-cell transcriptomes (SCTs). Excluding SCTs with <200 genes, 500 UMIs, or >10% mitochondrial genes resulted in 7,506 SCTs that were further analyzed. This subset had median counts of 2593 UMIs (min = 501, max = 17787, IQR = 3986) and 349 genes (min = 302, max = 5276, IQR = 1733) per cell (Figure S1C). Subsequent analysis of gene expression patterns by principal components (PC) analysis and shared-nearest-neighbors (Satija et al., 2015) resulted in 12 cellular clusters (Figure 1B, see Materials and Methods for details on sequential clustering). Major cell classes (i.e. neurons, astrocytes, microglia, etc…) within these clusters were identified by expression of cell-type specific gene combinations that were extensive cross-referenced with published datasets (Saunders et al., 2018; Zeisel et al., 2018) (Figure 1B-C). In contrast to other species (Pandey et al., 2018), no major transcriptional differences were observed (Figure S1A-B) across left and right hemispheres; therefore, cells from both hemispheres were pooled for analysis. The majority of the cells in the dataset were neurons (53%) and we focused our analysis on these clusters for the remainder of the study.

**Figure 1:**
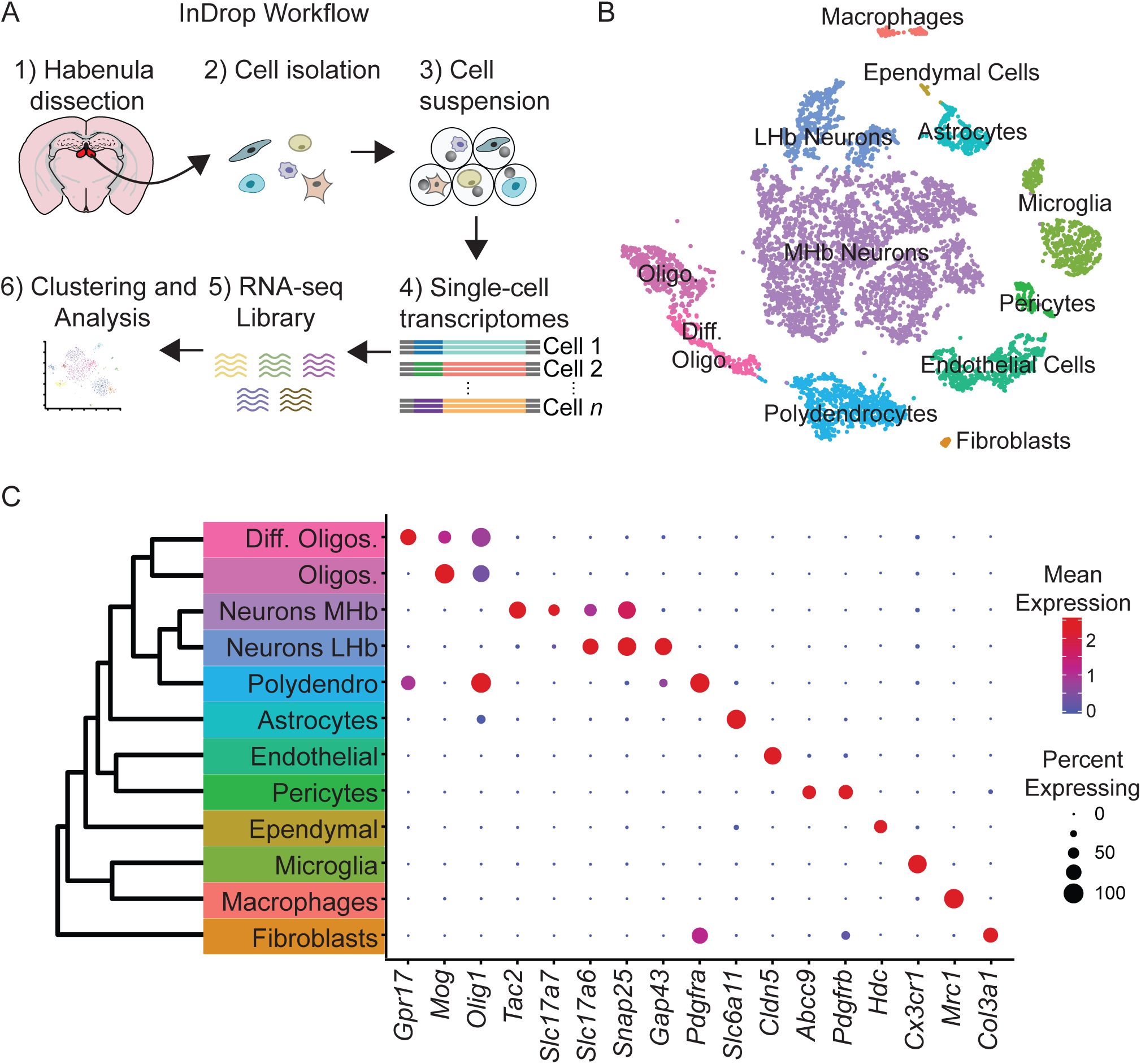
High-throughput single cell transcriptomic profiling of the habenula. **(A)** Schematic for scRNA-seq using the inDrop platform. Tissue containing the habenula was microdissected from acute coronal brain slices prepared from adult mice (1). Tissue chunks were digested in a cocktail of proteases and followed by trituration and filtration to obtain a cell suspension (2). Single cells were encapsulated using a droplet-based microfluidic device (3) for cell barcoding and mRNA capture (4). RNA sequencing (5) and bioinformatics analysis followed (6). **(B)** t-SNE plot of the processed dataset containing 7,506 cells from 6 animals. Cells are color-coded according to the cluster labels shown in (C). **(C)** Left: Dendrogram with cell class labels corresponding to clusters shown in (B). Right: Dot plot displaying expression of example enriched genes used to identify each major cell class. The color of each dot (blue to red) indicates the relative log-scaled expression of each gene whereas the dot size indicates the fraction of cells expressing the gene.

Neurons (*n* = 3,930 cells), identified by expression of genes required for chemical synaptic transmission such as *Snap25*, *Syp*, and *Syt4*, clustered into 2 main classes (Figure 1B-C). We examined if these 2 neuronal clusters could be spatially distinguished using digital *in situ* hybridization (ISH) analysis (Allen Brain Atlas, (Lein et al., 2007)) of differentially expressed genes (Finak et al., 2015). The larger cluster of neurons (n = 3,370 cells) expressed *Tac2* and corresponds to the MHb (Figure 2), whereas the smaller cluster (n = 560 cells) expressed *Gap43* and corresponds to the LHb (Figure 3).

**Figure 2:**
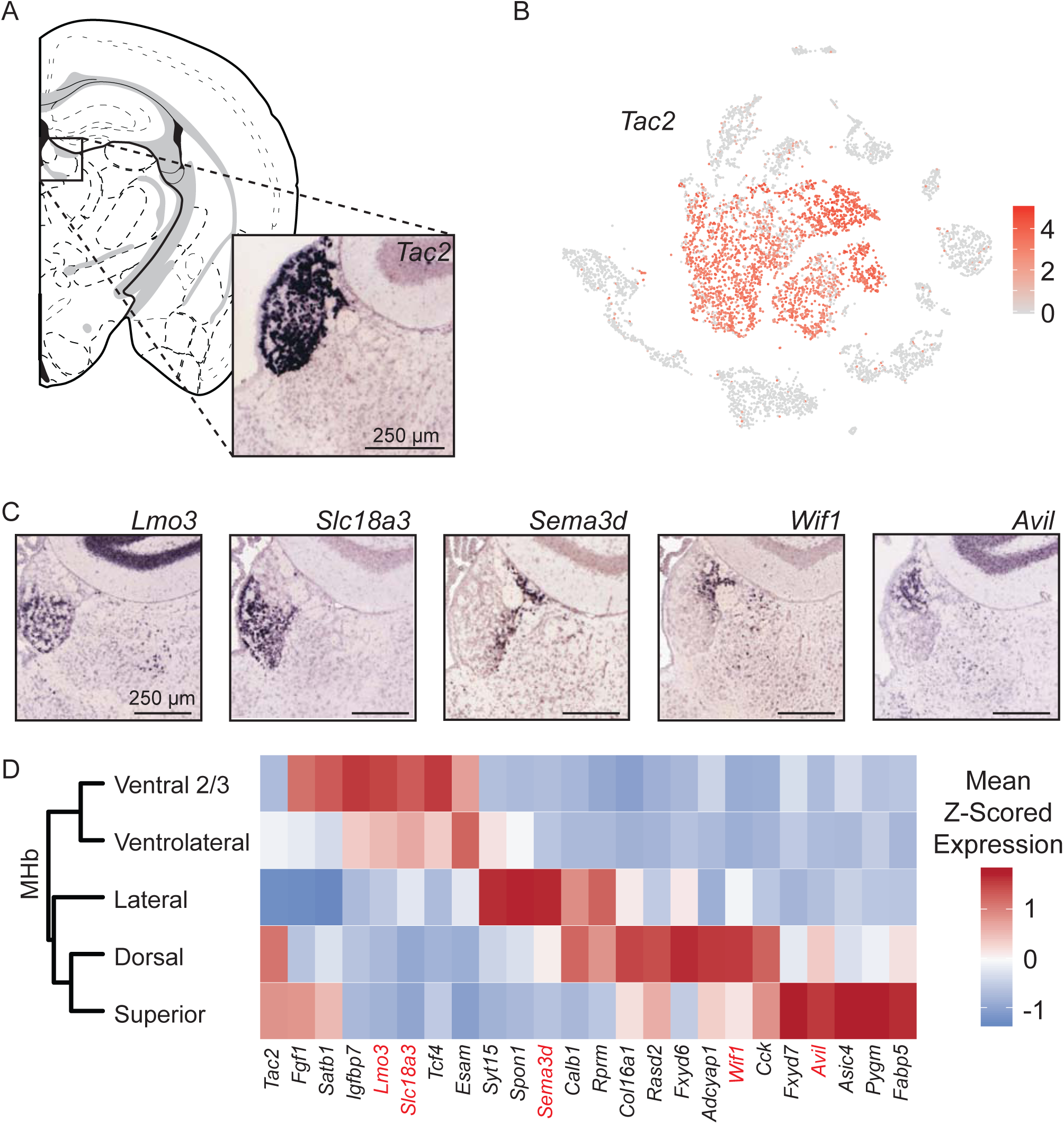
MHb neuron subtypes can be distinguished transcriptionally. **(A)** Location of MHb and ISH of *Tac2* expression from the Allen Institute Database. *Tac2* expression is restricted to cells in the MHb in this region. **(B)** *Tac2* serves as an excellent marker for MHb neurons in the dataset of SCTs (Scale on right shows normalized (log) gene expression.) **(C)** Sample ISH images from the Allen Institute Database showing selected differentially expressed genes for distinct transcriptionally defined neuronal subtypes in MHb. **(D)** Left: Dendrogram with MHb subtype labels corresponding to clusters shown in (Figure S2C). Right: Heatmap showing the relative expression (mean z-scored) of selected genes that are enriched in each MHb neuron subtype. Spatial distributions of enriched genes highlighted in (C) are labeled in red.

**Figure 3:**
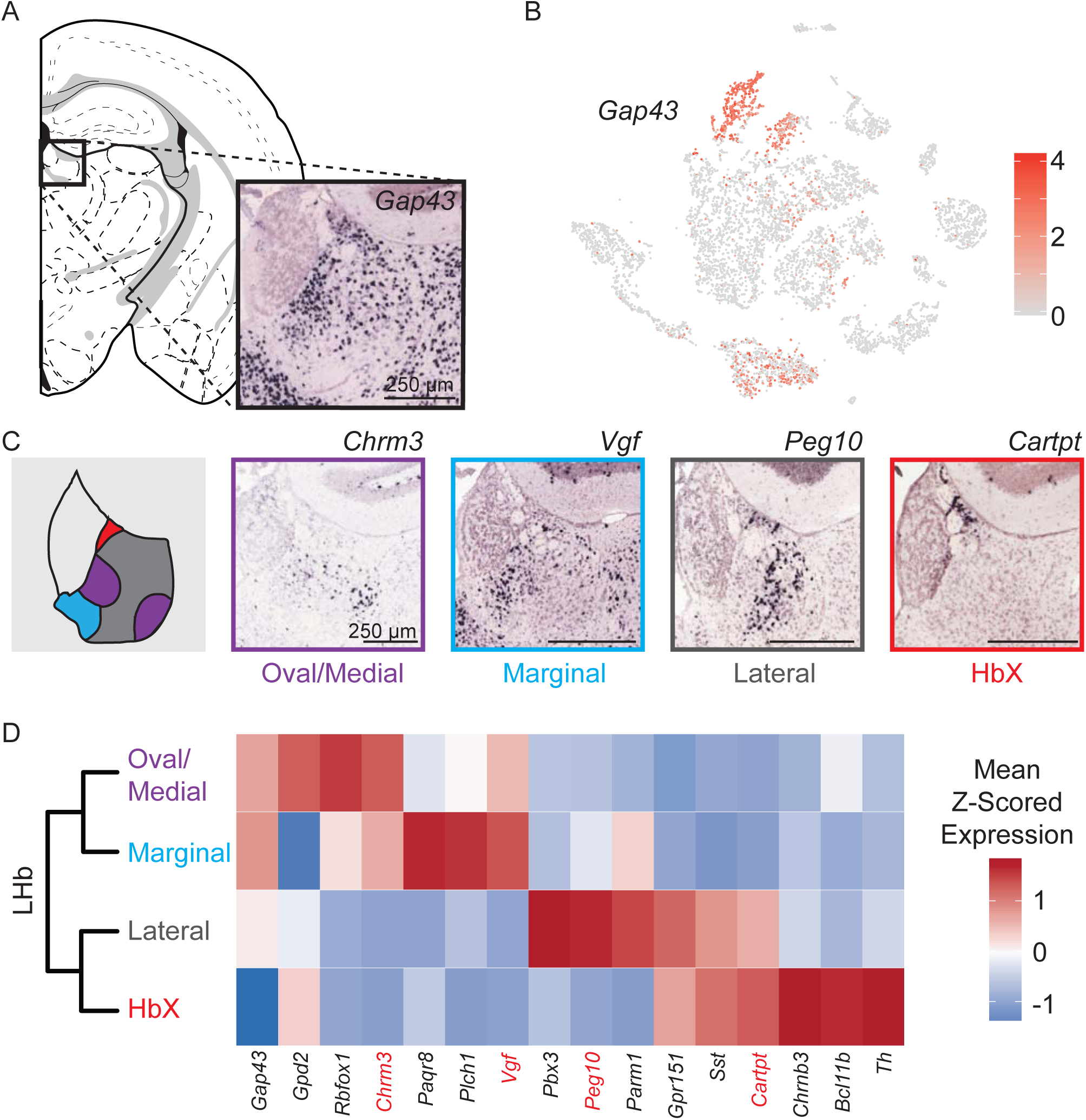
Characterization of genes differentially expressed between LHb neuron subtypes. **(A)** Location of LHb clusters and ISH *Gap43* expression from the Allen Institute Database. *Gap43* is highly expressed in neurons of the LHb and surrounding thalamus in this region, but excluded from MHb neurons. **(B)** *Gap43* serves as an excellent marker for LHb neurons in the dataset of single cell transcriptomes (Scale on right shows normalized (log) gene expression.) **(C)** Left: Illustration showing patterns of gene expression observed for genes shown for sample ISH to the right. Right: Sample ISH images from the Allen Institute Database showing selected differentially expressed genes for distinct transcriptionally defined neuronal subtypes in LHb. **(D)** Left: Dendrogram with LHb neuron labels corresponding spatial locations of differentially expressed genes within the LHb. Right: Heatmap showing the relative expression of selected genes that are enriched in each LHb neuron subtype. Spatial distributions of enriched genes highlighted in (C) are labeled in red.

### Differential gene expression reveals the spatial organization of MHb neuron subtypes

Analysis of MHb neurons revealed that they could be divided into 8 clusters (Figure S2A). However, 3 clusters were clearly distinguished by high expression of activity-dependent genes (ADGs) (Figure S2B), suggesting that they might simply reflect neurons of other clusters that had been recently strongly activated. Indeed, regressing out the PC containing a large number of ADGs (Figure S2E-F) caused these 3 high ADGs clusters to merge with other MHb clusters (Figure S2C-D), leaving 5 distinct subtypes of MHb neurons.

We constructed a cluster dendrogram using the averaged cluster gene expression to examine the transcriptional differences between these subtypes (Figure 2D). In general, subtypes of MHb neurons were divided by genes that were involved in the synthesis and packaging of different neurotransmitters and neuropeptides. All MHb neurons expressed high levels of *Slc17a6* and *Slc17a7*, the genes encoding vesicular glutamate transporters 1 and 2, and *Tac2* suggesting that all MHb neurons are glutamatergic and produce the neuropeptide Neurokinin B (Figure 2D, S4A). Two of the five clusters also expressed *Slc18a3* and *Chat* (not shown), the vesicular transporter and biosynthetic enzyme for acetylcholine, respectively, indicating that these neurons may co-release glutamate and acetylcholine (Figure S4A), as has been described in several studies (Ren et al., 2011; Soria-Gómez et al., 2015). Interestingly, no MHb neurons expressed significant levels of the canonical GABA handling genes *Slc32a1, Gad1*, or *Slc18a2* (although *Gad2* was expressed at low levels in all subtypes); therefore, they are unlikely to release GABA.

To examine the spatial distribution of MHb neuron subtypes we cross referenced their differentially expressed genes (DEGs) with the Allen Mouse Brain Atlas of ISH hybridization data (Table 3) (Lein et al., 2007). Generally, we found that individual DEGs for particular MHb subtypes consistently mapped onto discrete regions in the MHb (Figure 2C, S3). Also, DEGs for MHb neurons were rarely DEGs for LHb neurons (Figure S4A). This permitted classification of transcriptionally defined MHb subtypes to particular subregions of MHb (Figure 2D, S7). MHb neurons divided along the dorsal/ventral axis with a third lateral (enriched for genes *Sema3d, Calb1*, and *Spon1*) subtype (Figure 2C-D, S3C-E). Ventral groups could be further subdivided into two distinct subtypes, the “ventral two thirds” of the MHb (enriched for *Lmo3*) and the “ventrolateral” MHb (enriched for genes *Esam* and *Slc18a3*). Gene expression patterns indicated that it is possible that neurons from these two subtypes were partially intermingled and did not form a defined border (Figure 2C, S3A-B). The rest of the MHb could be subdivided into the “dorsal” (enriched for genes *Col16a1, Wif1*, and *Adcyap1*) and “superior” (enriched for genes *Cck* and *Avil*) subtypes. These two groups split along a medial/lateral axis with the “dorsal” being more laterally located than the “superior” (Figure 2C-D, S3F-I) (Wagner et al., 2016, 2014).

### Genetic distinction of four LHb neuron subtypes

*Gap43* is highly expressed in the LHb and along with several other genes (*Htr2c*, *Pcdh10, Gabra1*, and *Syn2*) distinguishes neurons of this region from those of neighboring MHb (Figure 3A-B, S4A). Unlike for MHb neurons, we did not detect significant elevation of ADGs in LHb neurons. We found 4 distinct clusters of neurons in LHb which, again unlike MHb, did not have distinct expression profiles of genes involved in the synthesis and packaging of different typical fast neurotransmitters (e.g. glutamate, GABA, acetylcholine) – all LHb neurons expressed high levels of *Slc17a6* and very low levels of *Slc32a1*, suggesting that they are glutamatergic.

Subdivisions of the lateral habenula based on topographic, morphological and cytochemical criteria have been described in rat (Andres et al., 1999) and mouse (Wagner et al., 2016, 2014) and are used here to describe the patterns of DEGs extracted from our single-cell dataset (see terms in quotes below). We examined the spatial distribution of LHb neuron subtypes by cross referencing their differentially expressed genes (DEGs) with the Allen Mouse Brain Atlas of ISH data (Lein et al., 2007). We found that DEGs showed 4 distinct, but consistent patterns that aligned with their subclusters (Figure 3C-D, S8). These consisted of 1) a cluster that showed high expression of DEGs in both the “lateral oval” and “central medial” subdivision, we named this the *oval/medial* subdivision; 2) a cluster that showed high expression of DEGs in the “marginal subdivision of the medial division of the LHb”, we called this the *marginal* subdivision; 3) a cluster that showed high expression of DEGs in the “lateral” subdivision (but avoiding expression in the “lateral oval”), we also called this the *lateral* subdivision; and 4) a cluster that showed high expression of DEGs in the subdivision defined as “HbX” lying on the dorsal border between MHb and LHb, we also refer to this as the *HbX* subdivision (Figure 3C) (Wagner et al., 2016, 2014). Interestingly, the *HbX* region is more closely related in its gene expression to other LHb clusters than to any clusters in the MHb; therefore it is more similar to LHb neurons than previously recognized (Figure 3D) (Wagner et al., 2016).

We performed multiplexed FISH to confirm that the 4 transcriptionally-defined clusters of LHb neurons were distinct and anatomically organized within the LHb. We chose 4 highly expressed DEGs (*Chrm3, Vgf, Gpr151*, and *Sst*) and examined gene expression levels in individual neurons (Figure 4). As predicted by the single cell sequencing, the chosen genes generally expressed in different cells, confirming that they defined molecularly distinct neuronal subtypes (Figure 4). An exception to this general rule, but consistent with the predictions of single cell sequencing, individual neurons in the HbX expressed both *Sst* and *Gpr151* (Figure 4D). Additionally, when strongly expressed, *Chrm3* and *Vgf* were found in different cells, but they were occasionally co-expressed in neurons that had relatively low levels of both genes (Figure 4A).

**Figure 4:**
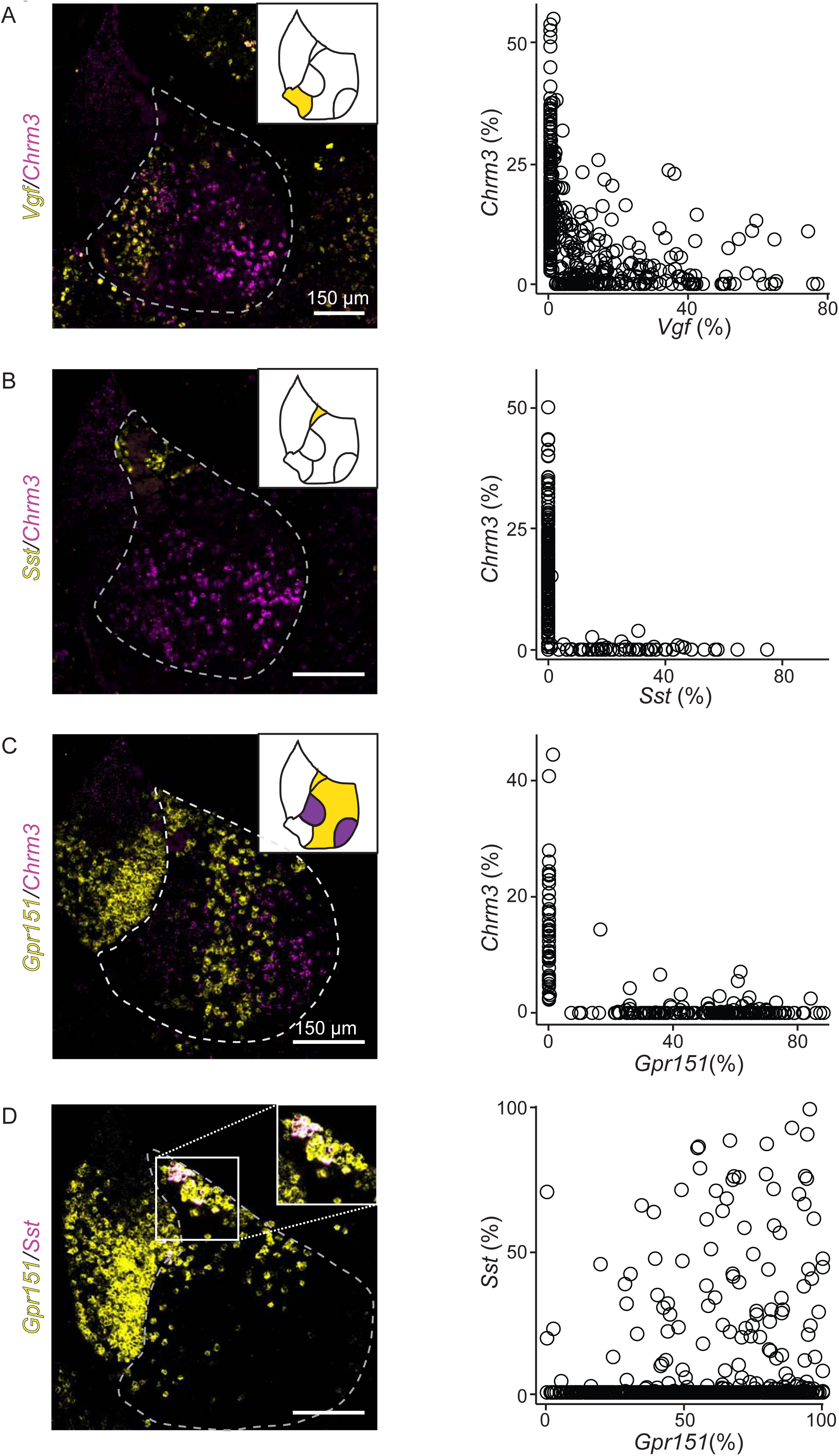
FISH confirms that differentially expressed genes from LHb subclusters are nonoverlapping and confined to specific spatial locations of LHb. **(A)** Left: Sample FISH of two differentially expressed LHb genes (*Vgf* (yellow) and *Chrm3 (*magenta)), with distinct spatial profiles (LHb outlined with gray dashed line). Right: Quantification of fluorescence coverage of single cells for FISH of *Vgf* and *Chrm3* in LHb neurons (n= 444 cells, 3 mice). **(B)** Left: Sample FISH of two differentially expressed LHb genes (*Sst* (yellow) and *Chrm3 (*magenta)), with distinct spatial profiles. Right: Quantification of fluorescence coverage of single cells for FISH of *Sst* and *Chrm3* in LHb neurons (n= 252 cells, 3 mice). **(C)** Sample FISH of two differentially expressed LHb genes (*Gpr151* (yellow) and *Chrm3 (*magenta)), with distinct spatial profiles (illustrated in upper right inset), LHb outlined in gray dashed line. **(F)** Quantification of fluorescence coverage of single cells for FISH of *Gpr151* and *Chrm3* in LHb neurons (n= 240 cells, 3 mice). **(D)** Left: Sample FISH of two differentially expressed LHb genes (*Sst* (yellow) and *Gpr151 (*magenta)), with distinct spatial profiles. Right: Quantification of fluorescence coverage of single cells for FISH of *Sst* and *Gpr151* in LHb neurons (n= 112 cells, 3 mice).

The chosen genes are largely expressed in non-overlapping patterns at the macroscopic level, confirming the organization of LHb into molecularly-defined subregions (Figure 4, S7). Nevertheless, cells from a subtype did intermingle with cells of another group and sharply defined borders between LHb subregions were not observed (e.g. Figure 4C). Therefore, diagrams of gene expression (Figure 4) illustrate where gene expression is greatest or where cells expressing the gene are most numerous and not that gene expression is perfectly restricted to a particular subregion.

### LHb neuron subtypes differentially target VTA GABAergic and dopaminergic neurons

The LHb projects via the fasciculus retroflexus to the ventral tegmental area (VTA), rostromedial tegmental area (RMTg), and median/dorsal raphe (Herkenham and Nauta, 1977). The VTA consists of a large and diverse population of dopamine neurons, as well as smaller populations of purely GABAergic, purely glutamatergic, and GABA/glutamate coreleasing neurons. Both GABAergic and dopaminergic VTA neurons receive input from the LHb (Beier et al., 2015; Lammel et al., 2012; Morales and Margolis, 2017; Watabe-Uchida et al., 2012) but it is unknown if these arise from molecularly distinct LHb neurons. We tested if there was connectivity specificity between LHb and VTA neuronal subtypes using rabies virus based monosynaptic retrograde tracing (Wickersham et al., 2007). To examine LHb input to VTA GABAergic neurons we injected Cre-dependent TVA-mCherry into the VTA of a VGAT-IRES-Cre mouse to restrict initial rabies virus infection to GABAergic neurons. We also coinjected a Cre-dependent AAV encoding the rabies glycoprotein (RVG) to allow for retrograde monosynaptic transfer of G-deleted, pseudotyped, rabies virus (EnvA-RbV-GFP). As only neurons with Cre will express RVG, GFP-labeled neurons in other regions are putatively presynaptic to GFP+/RVG+ VTA neurons (see Figure S6E for controls for specificity of EnvA-RbV-GFP infection). FISH in the VTA revealed that ∼30% of “starter cells” (neurons that were GFP+ and Cre+), coexpressed *Slc17a6* indicating they are likely GABA/glutamate coreleasing neurons (Figure S6C) (Root et al., 2014). The majority of the remaining 70% of “starter cells” are purely GABAergic (Figure S6C-D).

Using FISH we found retrogradely labeled neurons, marked by expression of *RbV-N* mRNA, in all four LHb subtypes (identified using enriched genes *Chrm3, Vgf, Gpr151*, and *Sst*) (Figure 5C-D, S5A). The majority of retrogradely labeled LHb neurons were found in the lateral and oval/medial subtypes in roughly equal proportions (mean±SEM: 48±0.5% *Gpr151+* and 41±0.6% *Chrm3+*, respectively) (Figure 5D). A much smaller proportion was found in the marginal subtype (10±1% *Vgf+*), and very few HbX neurons (2±1% *Sst+*) were retrogradely labeled.

**Figure 5:**
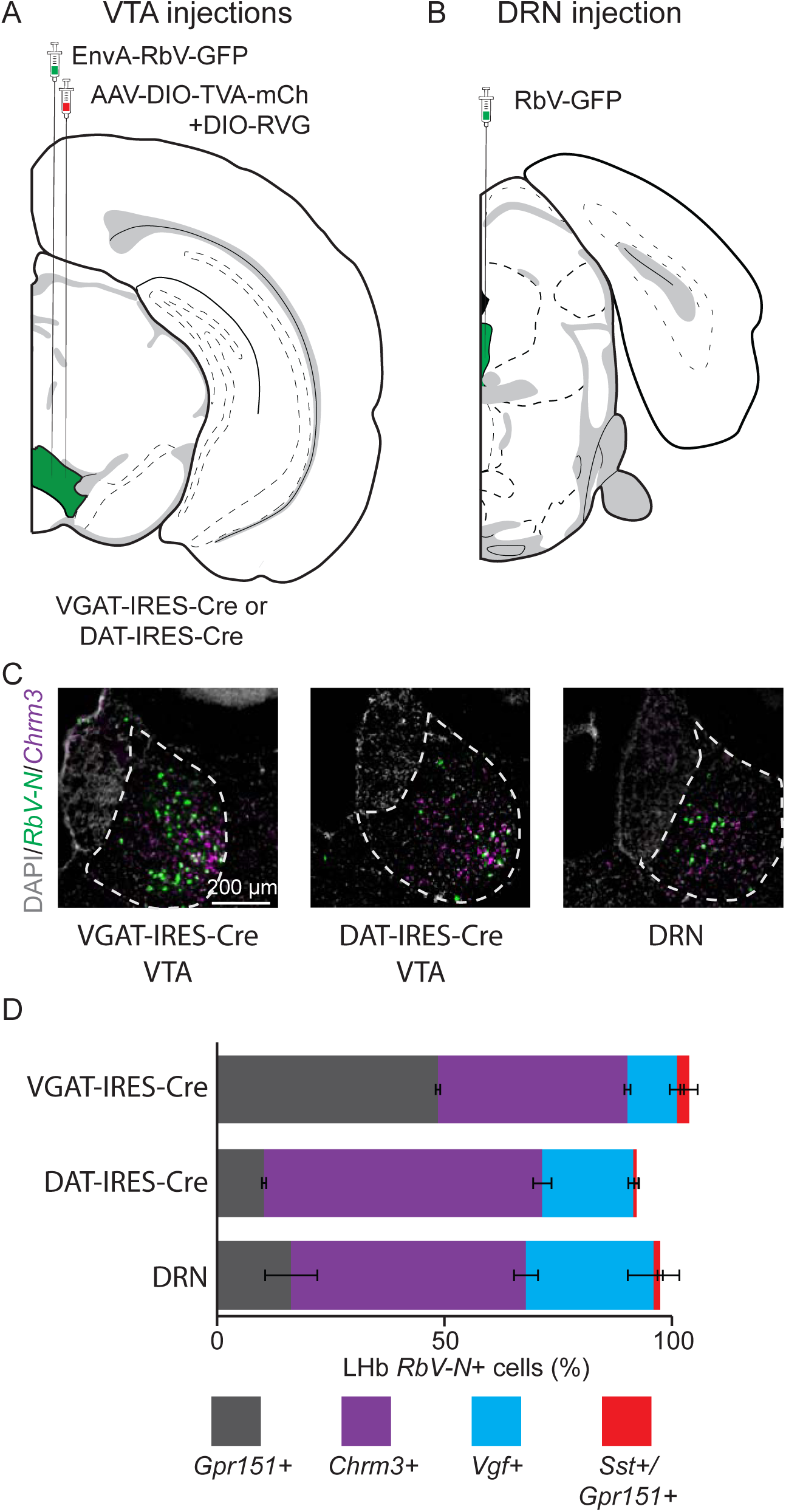
Distinct LHb neuron subtypes prefer different downstream targets, but all subtypes target both VTA and DRN. **(A)** Location of sites for AAV helper viruses (AAV-FLEX-TVA-mCh and AAV-FLEX-RVG) and pseudotyped rabies virus (EnvA-RbV-GFP) injection into VTA. **(B)** Location of non-pseudotyped rabies virus (RbV-GFP) injection into DRN. **(C)** Sample habenula FISH images for *RbV-N* and *Chrm3* following viral injection into either VTA or DRN. **(D)** Quantification of the proportion of *RbV-N* labeled neurons that overlapped with the enriched genes for distinct LHb neuron subtypes (VGAT-IRES-Cre n= 1430 cells/ 4 mice, DAT-IRES-Cre n= 549/ 3 mice, DRN n= 465/ 3 mice). Filled rectangles are the mean and error bars are ±SEM, see Table 6 for statistical comparisons.

To examine LHb input to VTA dopaminergic neurons we performed monosynaptic retrograde tracing using the same series of viral injections in DAT-IRES-Cre mice (Figure 5A). FISH in the VTA revealed that ∼91% of “starter cells” (neurons that were GFP+ and Cre+), coexpressed *Slc6a3* (dopamine transporter, DAT) indicating they are dopaminergic neurons (Figure S6D) (Morales and Margolis, 2017; Tritsch et al., 2012). The remaining 9% of starter cells express varying levels of *Slc32a1* (VGAT) indicating low levels of starter cell overlap with the experiments done in the VGAT-IRES-Cre line (Figure S6D). We performed FISH on retrogradely labeled neurons in the LHb and found that a much larger proportion of retrogradely labeled neurons in the oval/medial and marginal subtypes (61±2% *Chrm3+* and 20±1% *Vgf+*, respectively) in the DAT-IRES-Cre than the VGAT-IRES-Cre line (Figure 5C-D, S5B). Consequently, a smaller proportion of neurons in the lateral subtype were labeled (10±0.4% *Gpr151+*) and almost no neurons in the HbX subregion were labeled (0.7±0.5% *Sst+*) (Figure 5D, S5B). Together these data suggest that both VTA GABAergic and dopaminergic neurons can receive input from all 4 subtypes of LHb neuron. However, VTA dopaminergic neurons receive the largest proportion of their LHb input from the oval/medial LHb subtype, whereas VTA GABAergic neurons receive equal levels of input from both the oval/medial and lateral LHb subtypes (Table 6).

### All LHb neuron subtypes project to the DRN in proportions similar to VTA dopamine neurons

LHb neurons heavily innervate the dorsal and median raphe nuclei (DRN/MRN) and modulate serotonergic output throughout the brain (Zhao et al., 2015; Zhou et al., 2017). To examine the LHb subtypes that project to the DRN, we injected a non-pseudotyped rabies virus (RbV-GFP) into this area and performed FISH in the LHb for subtype enriched genes (Figure 5B-C). Similar to dopaminergic VTA neurons, the DRN received the largest proportion of its LHb input from the oval/medial subtype (51±2% *Chrm3+*) (Figure 5C-D, S5C). The DRN also received minor inputs from the lateral (16±5% *Gpr151+*), marginal (28±5% *Vgf+*), and HbX (1±0.5% *Sst+*) regions (Figure 5D, S5C). Overall, the proportions of input to the DRN arising from different LHb subtypes were more similar to those to VTA dopamine neurons than those to VTA GABA neurons (Table 6).

## DISCUSSION

We performed transcriptional and anatomical analyses of the habenula, a crucial circuit node that modifies brain-wide dopamine and serotonin levels through its connections to the VTA and DRN (Proulx et al., 2014; Tian and Uchida, 2015; Zhao et al., 2015). Using large-scale single cell transcriptional profiling, we classify MHb and LHb neurons into five and four major neurons types, respectively, and show that each class has a distinct gene expression pattern. The four LHb populations were confirmed to be non-overlapping at the single-cell level by FISH. Monosynaptic retrograde tracing revealed that GABAergic VTA neurons receive equal input from the oval/medial and lateral LHb neuronal subtypes, whereas dopaminergic VTA neurons receive input primarily from the oval/medial LHb subtype. Finally, neurons of the DRN receive input from the LHb in roughly similar proportions to dopaminergic VTA neurons.

### Anatomical distribution of MHb neuronal subtypes

Recent studies have identified and delineated the subnuclei of the mouse MHb using morphological, topographic and cytochemical criteria (Wagner et al., 2016, 2014). Using single-cell transcriptional profiling, we show that MHb neurons can be categorized into subtypes based on differential gene expression. Furthermore, the spatial distribution of these transcripts allowed us to ascribe an anatomical location to each subtype. The anatomical location of these subtypes largely agree with previously defined MHb subnuclei and we have used the same nomenclature when possible (Figure S7) (Wagner et al., 2016).

The two ventral subtypes of the MHb coexpressed transcripts for glutamate and ACh neurotransmission (Figure S4). Our data suggest these two ventral subtypes can be differentially targeted with intersectional approaches, as genes such as *Lmo3* and *Esam* are preferentially expressed in one subtype (Figure 2D, S4). Previous studies indicate that MHb neurons that release glutamate and ACh target the medial interpeduncular nucleus (IPN) (Ren et al., 2011) and are involved in the formation of aversive memories (Soria-Gómez et al., 2015). However, whether one or both of the transcriptionally-defined subtypes are involved in this process is unknown.

Additionally, cholinergic transmission in MHb has also been implicated in nicotine addiction as MHb neurons not only release ACh, but express an array of nicotinic acetylcholine receptor subunits (nAChRs, such as *Chrna3* and *Chrnb3;* Figure S3 and Table 4) (Fowler et al., 2011; Shih et al., 2014). Similar to its involvement in aversive memories, MHb likely plays an important role in mediating the unpleasant symptoms associated with nicotine withdrawal (Zhao-Shea et al., 2013). Our data provide a comprehensive view of all nAChR and mAChR transcripts expressed in both MHb and LHb providing a resource for the development of new therapeutic targets for the treatment of addiction (Table 4) (D’Souza, 2016; Zuo et al., 2016).

Few studies have examined the function of the dorsal (enriched for genes *Col16a1, Wif1*, and *Adcyap1*) and superior (enriched for genes *Cck*, and *Avil*) MHb. These neurons were known to express high levels of *Tac1* (the gene that produces the neuropeptide substance P), consistent with our single-cell sequencing data (Figure S3J, Table 4) and target the lateral IPN (Hsu et al., 2016). Their activation may be reinforcing (Hsu et al., 2014), but detailed analysis of their function and neurotransmitter release has not been examined.

### LHb neuronal subtypes

We referenced recent studies on LHb subnuclei to create a map (Figure S7) of LHb based on DEGs extracted from single-cell transcriptional profiling (Wagner et al., 2016, 2014). Overall, our map largely agrees with previous work and adds many key observations into the organization and cellular and molecular diversity of the LHb. In addition to providing multiple genetic handles that can be used in future studies to target LHb neuron subtypes, our study reveals the a wide range of GPCRs (such as *Htr2c, Htr5b, Sstr2, Gpr151*; see Table 4) expressed in LHb neurons that could be targeted for treatment of diseases known to effect LHb function such as depression, anxiety, and addiction (Lecca et al., 2014; Proulx et al., 2014). In contrast to some reports (Zhang et al., 2018), we did not find evidence of GABAergic neurons in the LHb (or MHb). Although *Gad2* and *Slc6a1*, which encode a GABA synthetic enzyme and GABA transporter, respectively, were present at low levels in all LHb clusters we did not find expression of *Slc32a1* or *Slc18a2*, which are required for vesicular loading of GABA (Table 4). This is in agreement with recently published results demonstrating that *Gad2* expression is a poor discriminator for inhibitory (GABAergic) neurons (Moffitt et al., 2018). Therefore, either LHb GABAergic cells are rare enough to be missed in the single cell transcriptomes, or the habenula is devoid of GABAergic neurons.

We used *Gpr151* expression to mark the lateral, and to a lesser extent HbX, regions of the LHb. The *Gpr151*+ neurons of the lateral LHb are the most well-studied neuronal subtype in the LHb and they receive major input from the lateral preoptic area, lateral hypothalamus, entopeduncular nucleus (EP), basal nucleus of the stria terminalis, and the nucleus of the diagonal band (Broms et al., 2017). These neurons also receive a minor input from the VTA, and are positioned to receive GABA/glutamate coreleasing input from both the EP and VTA (Root et al., 2014; Wallace et al., 2017). *Gpr151*+ axons, likely arriving from the LHb, heavily innervate the RMTg and central and median raphe nucleus, but not the VTA (Broms et al., 2015). Our retrograde tracing studies from VTA neurons show that *Gpr151+* neurons tend to avoid dopaminergic VTA neurons, but heavily innervated the intermingled GABAergic neurons in this brain region (Figure 5). VTA GABAergic interneurons are functionally similar to inhibitory RMTg neurons (both populations inhibit VTA dopaminergic neurons (Cohen et al., 2012; Ji and Shepard, 2007)), consistent with our results that both are innervated by lateral LHb. Furthermore, the lateral LHb is likely the major LHb subtype to translate aversive signals to VTA GABAergic neurons which become active following reward omission due to increased LHb input (Tian and Uchida, 2015). Overall, these cells are positioned to translate signals arriving from EP to downstream midbrain structures involved in both dopamine and serotonin signaling.

The oval/medial subregion of the LHb expresses high levels of *Chrm3*. Our previous studies indicate that this subtype is positioned to receive purely glutamatergic input from *Pvalb+/Slc16a7+* EP neurons that specifically target the lateral oval nucleus of the LHb (Wallace et al., 2017). Additionally, GABA/glutamate coreleasing EP neurons target the lateral oval (as well as the neighboring *Gpr151+* lateral LHb) providing overlapping, but differential EP input to this subregion (Wallace et al., 2017). Electrophysiological analysis has also shown preferential input from EP to the lateral LHb, specifically, to the neurons that project to RMTg (Meye et al., 2016).

The LHb primarily targets meso-prefrontal VTA dopamine neurons while avoiding other (mesolimbic and substantia nigra) dopamine neurons (Lammel et al., 2012). We were surprised to find that the majority of LHb neurons that projected to VTA dopamine neurons expressed *Chrm3*, a gene enriched in the oval/medial subregion, as previous retrograde tracing studies suggest that the marginal portion of the LHb projects heavily to the VTA ((Meye et al., 2016), but also see (Petzel et al., 2017; Quina et al., 2014)). Additional studies suggest that LHb neurons that target the VTA dopamine neurons may be distinct from those that target the RMTg; therefore, two populations of *Chrm3+* LHb neurons with different synaptic targets may exist (Lammel et al., 2012; Li et al., 2011; Maroteaux and Mameli, 2012). Additional genetic heterogeneity between *Chrm3+* oval and medial subdivisions could be further resolved with higher resolution sequencing methods (Bakken et al., 2018; Tasic et al., 2018). Nevertheless, our data show that VTA dopamine, and VTA GABAergic neurons are positioned to receive quite different synaptic input from the LHb due to their differential targeting by LHb neuronal subtypes.

*Vgf* expression was enriched in the marginal subtype of the LHb. Our retrograde tracing studies revealed that this subregion projects most heavily to the DRN, similar to other studies showing labeling of the medial half of the LHb following injections of retrograde tracers into the raphe nucleus (Quina et al., 2014). Interestingly, this region also appears to receive dense input from serotonergic neurons of the raphe nuclei (Huang et al., 2019), and express *Htr2c* as well as several other serotonin receptors (See Table 4). We expect that this subregion also projects heavily to the lateral dorsal tegmental nucleus (LDTg) and posterior hypothalamic area (PH), as retrograde injections into these areas exclusively label the medial half of the LHb (Quina et al., 2014).

### Summary

Progress in defining a function for the habenula has been hindered by incomplete understanding of its constituent cell-types and subregions. This study provides a comprehensive description of the neuronal classes in the lateral and medial habenula based on single-cell transcriptional profiling, FISH, and cell type specific monosynaptic retrograde tracing (Figure S7). Future studies will improve our understanding of the function of these habenula cell types by employing current optogenetic, chemogenetic, and electrophysiological approaches for precise control and monitoring of individual habenular populations.

## MATERIALS AND METHODS

### Mice

The following mouse strains/lines were used in this study: C57BL/6J (The Jackson Laboratory, Stock # 000664), *VGAT-IRES-Cre* (The Jackson Laboratory, Stock # 016962), *DAT-IRES-Cre* (The Jackson Laboratory, Stock # 006660). Animals were kept on a 12:12 regular light/dark cycle under standard housing conditions. All procedures were performed in accordance with protocols approved by the Harvard Standing Committee on Animal Care following guidelines described in the U.S. National Institutes of Health Guide for the Care and Use of Laboratory Animals.

### Adeno-Associated Viruses (AAVs)

Recombinant AAVs used for retrograde tracing experiments (AAV2/9-CAG-FLEX-TCB-mCherry, AAV2/9-CAG-FLEX-RVG) were commercially obtained from the Boston Children’s Hospital Viral Core (Addgene # 48332 and 48333, respectively). Virus aliquots were stored at −80 °C, and were injected at a concentration of approximately 10^11^ or 10^12^ GC/ml, respectively.

### Rabies Viruses

Rabies viruses used for retrograde tracing (B19G-SADΔG-EGFP) were generated in-house (Wickersham et al., 2010). Virions were amplified from existing stocks in three rounds of low-MOI passaging through BHK-B19G cells by transfer of filtered supernatant, with 3 to 4 days between passages. Cells were grown at 35 °C and 5% CO_2_ in DMEM with GlutaMAX (Thermo Scientific, #10569010) supplemented with 5% heat-inactivated FBS (Thermo Scientific #10082147) and antibiotic-antimycotic (Thermo Scientific #15240-062). Virions were concentrated from media from dishes containing virion-generating cells by first collecting and incubating with benzonase nuclease (1:1000, Millipore #70664) at 37°C for 30 min, followed by filtration through a 0.22 µm PES filter. The filtered supernatant was transferred to ultracentrifuge tubes (Beckman Coulter #344058) with 2 ml of 20% sucrose in dPBS cushion and ultracentrifugated at 20,000 RPM (Beckman Coulter SW 32 Ti rotor) at 4°C for 2 hours. The supernatant was discarded and the pellet was resuspended in dPBS for 6 hours on an orbital shaker at 4 °C before aliquots were prepared and frozen for long-term storage at −80 °C. Unpseudotyped rabies virus titers were estimated based on a serial dilution method counting infected HEK 293T cells, and quantified as infectious units per ml (IU/ml). Pseudotyped rabies virus (SAD B19 strain, EnvA-RbV-GFP, Addgene# 52487) was commercially obtained from the Janelia Viral Tools Facility stored at −80°C, and injected at a concentration of approximately 10^8^ IU/ml.

### Stereotaxic Surgeries

Adult mice were anesthetized with isoflurane (5%) and placed in a small animal stereotaxic frame (David Kopf Instruments). After exposing the skull under aseptic conditions, viruses were injected through a pulled glass pipette at a rate of 50 nl/min using a UMP3 microsyringe pump (World Precision Instruments). Pipettes were slowly withdrawn (< 100 µm/s) at least 10 min after the end of the infusion. Following wound closure, mice were placed in a cage with a heating pad until their activity was recovered before returning to their home cage. Mice were given pre- and post-operative subcutaneous ketoprofen (10mg/kg/day) as an analgesic, and monitored daily for at least 4 days post-surgery. Injection coordinates from Bregma for VTA were −3.135mm A/P, 0.4mm M/L, and 4.4mm D/V and for DRN were −6.077mm A/P, 0.1mm M/L, and − 3.33mm D/V at −40°. Injection volumes for specific anatomical regions and virus types were as follows VTA: 200 nL AAV (mix of helper viruses), 250 nL EnvA-RbV-GFP (21 days after injection of AAV), DRN: 300 nL of RbV-GFP. Animals injected with rabies virus were perfused 7 days after injection in a biosafety level 2 animal facility.

### Single Cell Dissociation and RNA Sequencing

8- to 10-week old C57BL/6J mice were pair-housed in a regular 12:12 light/dark cycle room prior to tissue collection. Mice were transcardially perfused with an ice-cold choline cutting solution (110 mM choline chloride, 25 mM sodium bicarbonate, 12 mM D-glucose, 11.6 mM sodium L-ascorbate, 10 mM HEPES, 7.5 mM magnesium chloride, 3.1 mM sodium pyruvate, 2.5 mM potassium chloride, 1.25 mM sodium phosphate monobasic, saturated with bubbling 95% oxygen/5% carbon dioxide, pH adjusted to 7.4 using sodium hydroxide). Brains were rapidly dissected out and sliced into 200 µm thick coronal sections on a vibratome (Leica Biosystems, VT1000) with a chilled cutting chamber filled with choline cutting solution. Coronal slices containing the habenula were then transferred to a chilled dissection dish containing a choline-based cutting solution for microdissection. Dissected tissue chunks were transferred to cold HBSS-based dissociation media (Thermo Fisher Scientific Cat. # 14170112, supplemented to final content concentrations: 138 mM sodium chloride, 11 mM D-glucose, 10 mM HEPES, 5.33 mM potassium chloride, 4.17 mM sodium bicarbonate, 2.12 mM magnesium chloride, 0.441 mM potassium phosphate monobasic, 0.338 mM sodium phosphate monobasic, saturated with bubbling 95% oxygen/5% carbon dioxide, pH adjusted to 7.35 using sodium hydroxide) and kept on ice until dissections were completed. Dissected tissue chunks for each sample were pooled for each hemisphere for the subsequent dissociation steps. Tissue chunks were first mixed with a digestion cocktail (dissociation media, supplemented to working concentrations: 20 U/ml papain, 0.05 mg/mL DNAse I) and incubated at 34 °C for 90 min with gentle rocking. The digestion was quenched by adding dissociation media supplemented with 0.2% BSA and 10 mg/ml ovomucoid inhibitor (Worthington Cat. # LK003128), and samples were kept chilled for the rest of the dissociation procedure. Digested tissue was collected by brief centrifugation (5 min, 300 *g*), re-suspended in dissociation media supplemented with 0.2% BSA, 1 mg/ml ovomucoid inhibitor, and 0.05 mg/mL DNAse I. Tissue chunks were then mechanically triturated using fine-tip plastic micropipette tips of progressively decreasing size. The triturated cell suspension was filtered through a 40 µm cell strainer (Corning 352340) and washed in two repeated centrifugation (5 min, 300 *g*) and re-suspension steps to remove debris before a final re-suspension in dissociation media containing 0.04% BSA and 15% OptiPrep (Sigma D1556). Cell density was calculated based on hemocytometer counts and adjusted to approximately 100,000 cells/ml. Single-cell encapsulation and RNA capture on the InDrop platform was performed at the Harvard Medical School ICCB Single Cell Core using v3 chemistry hydrogels based on previously described protocols (Zilionis et al., 2017). Suspensions were kept chilled until the cells were flowed into the microfluidic device. Libraries were prepared and indexed following the protocols referenced above, and sequencing-ready libraries were stored at −80 °C. Libraries from different samples were pooled and sequenced on an Illumina NextSeq 500 (High Output v2 kits).

### Sequencing Data Processing

NGS data was processed using previously a published pipeline in Python available at [https://github.com/indrops/indrops] (Klein et al., 2015). Briefly, reads were filtered by expected structure and sorted by the corresponding library index. Valid reads were then demultiplexed and sorted by cell barcodes. Cell barcodes containing fewer than 250 total reads were discarded, and remaining reads were aligned to a reference mouse transcriptome (Ensembl GRCm38 release 87) using Bowtie 1.1.1 (m = 200, n = 1, l = 15, e = 1000). Aligned reads were then quantified as UMI-filtered mapped read (UMIFM) counts. UMIFM counts and quantification metrics for each cell were combined into a single file sorted by library and exported as a gunzipped TSV file.

### Pre-Clustering Filtering and Normalization

Analysis of the processed NGS data was performed in R (version 3.4.4) using the Seurat package (version 2.3.4) (Butler et al., 2018; Satija et al., 2015). Cells with fewer than 500 UMIFM counts and 200 genes were removed. The expression data matrix (Genes × Cells) was filtered to retain genes with > 5 UMIFM counts, and then loaded into a Seurat object along with the library metadata for downstream processing. The percentage of mitochondrial transcripts for each cell (percent.mito) was calculated and added as metadata to the Seurat object. Cells in the object were further filtered using the following parameters: nUMI – min. 500, max. 18000; nGene – min. 200, max. 6000; percent.mito – min. −Inf, max. 0.1. Low quality libraries identified as outliers on scatter plots of quality control metrics (e.g. unusually low gradient on the nGene vs. nUMI) were also removed from the dataset. Filtered Seurat objects were then log-normalized at 10,000 transcripts per cell. Effects of latent variables (nUMI, percent.mito) were estimated and regressed out using a GLM (`ScaleData` function, `model.use = “linear”`), and the scaled and centered residuals were used for dimensionality reduction and clustering.

### Cell Clustering and Cluster Identification

Initial clustering was performed on the dataset using the first 20 PCs, and t-SNE was used only for data visualization. Clustering was run using the SNN-based `FindClusters` function using the SLM algorithm and 10 iterations. Clustering was performed at varying resolution values, and we chose a final value of 1.2 for the resolution parameter for this stage of clustering. Clusters were assigned preliminary identities based on expression of combinations of known enriched genes for major cell classes and types. The full list of enriched genes is provided in Table 2 and average expression of all genes in all clusters is provided in Table 1. Low quality cells were identified based on a combination of low gene/UMIFM counts and high levels of mitochondrial and nuclear transcripts (e.g. *Malat1*, *Meg3*, *Kcnq1ot1*) typically clustered together and were removed. Following assignment of preliminary identities, cells were divided into data subsets as separate Seurat objects (LHb neurons and MHb neurons) for further subclustering. The expression matrix for each data subset was further filtered to include only genes expressed by the cells in the subset (minimum cell threshold of 0.5% of cells in the subset). Subclustering was performed iteratively on each data subset to resolve additional cell types and subtypes. Briefly, clustering was run at high resolution, and the resulting clusters were ordered in a cluster dendrogram using the `BuildClusterTree` function in Seurat which uses cluster averaged PCs for calculating a PC distance matrix. Putative doublets/multiplets were identified based on expression of known enriched genes for different cell types not in the cell subset (e.g. neuronal and glial specific genes). Putative doublets tended to separate from other cells and cluster together, and these clusters were removed from the dataset. Cluster separation was evaluated using the `AssessNodes` function and inspection of differentially expressed genes at each node. Clusters with poor separation, based differential expression of mostly housekeeping genes, or activity dependent genes (see Figure S2) were merged to avoid over-separation of the data. The dendrogram was reconstructed after merging or removal of clusters, and the process of inspecting and merging or removing clusters was repeated until all resulting clusters could be distinguished based on a set of differentially expressed genes that we could validate separately. To calculate the “ADG Score” (Figure S2) we used the `AddModuleScore` function in Seurat using a list of ADGs that were highly expressed in some of the MHb clusters (*Fos, Fosb, Egr1, Junb, Nr4a1, Dusp18, Jun, Jund*).

### Differential Expression Tests

Tests for differential gene expression were performed using MAST (version 1.10.1) (Finak et al., 2015) through the `FindMarkersNode` function in Seurat (`logfc.threshold = 0.25`, `min.pct = 0.1`). Adjusted *P* values were corrected using the Bonferroni correction for multiple comparisons (*P* < 0.05).

### Fluorescence In-Situ Hybridization (FISH)

Mice were deeply anesthetized with isoflurane, decapitated, and their brains were quickly removed and frozen in tissue freezing medium on dry ice. Brains were cut on a cryostat (Leica CM 1950) into 30 µm sections, adhered to SuperFrost Plus slides (VWR), and immediately refrozen. Samples were fixed 4% paraformaldehyde and processed according to ACD RNAscope Fluorescent Multiplex Assay manual. Sections were incubated at room temperature for 30 seconds with DAPI, excess liquid was removed, and immediately coverslipped with ProLong antifade reagent (Molecular Probes). Antisense probes for *RbV-N, Gpr151, Sst, Chrm3, Vgf, Cre, Slc17a6, Slc32a1, and Slc6a3* were purchased from Advanced Cell Diagonstics (ACD, http://acdbio.com/). Sections were imaged at 1920 × 1440 pixels on a Keyence BZ-X710 fluorescence microscope using a 10X, 0.45 NA air Nikon Plan Apo objective. Individual imaging planes were overlaid and quantified for colocalization in ImageJ (NIH) and Matlab (Mathworks).

### Image Analysis

FISH images were analyzed for “fluorescence coverage (%),” meaning the proportion of fluorescent pixels to total pixels in a cellular ROI, using a custom macro in Image J and custom scripts in Matlab (Figure 4, S5, and S6). 5-10 images from at least 3 mice were analyzed for each condition. Cell ROIs were automatically determined based on fluorescence signals in all three channels (or by fluorescence in the *RbV-N* channel for rabies tracing experiments), and manually adjusted prior to analysis to ensure that all cell ROIs reflected individual cells and not clusters. After background subtraction (the signal outside of cell ROIs) and application of a fluorescence threshold (Renyi Entropy), the amount of fluorescent pixels in each optical channel was counted within the cellular ROI. All images compared underwent identical thresholding and no other manipulations were made. These data were used to generate X-Y plots displaying the percent coverage for each channel per cell (Figure 4, S5, and S6).

## ACKNOWLEDGEMENTS

The authors would like to thank Sarah Melzer and Adam Granger for assistance with FISH analysis; James Levasseur for animal husbandry and genotyping; L. Worth for administrative assistance; HMS ICCB Single Cell Core for assistance with scRNA-seq experiments on the InDrop platform; The Bauer Core Facility at Harvard University for sequencing support and the members of the Sabatini Lab for their helpful discussions and advice. Starting materials for generating nonpseudotyped rabies virus is a generous gift from B.K. Lim (UCSD). This work was supported by the Howard Hughes Medical Institute (B.L.S.), NIH, National Institute of Neurological Disease and Stroke (K99 NS105883 to M.L.W and NS103226 to B.L.S.).

## SUPPLEMENTARY INFORMATION

**Figure S1 (related to Figure 1): Comparison of cell type composition across hemispheres and gene diversity, mitochondrial genes, and UMIs across cell types.**

**(A)** t-SNE plot of the dataset with cells color-coded by the hemisphere from which the sample was acquired. **(B)** Bar plots showing the percentage of cells in each hemisphere that are categorized into each of the 12 major cell types. **(C)** Violin plots of the number of genes (top), unique molecular identifiers (UMIs, middle), and percentage of mitochondrial genes (bottom) for each of the 12 cell types. Each point represents a single cell and filled area is a probability distribution of all the cells in that category.

**Figure S2 (related to Figure 2): Subclustering of MHb neurons before and after subtraction of heterogeneity introduced by elevated expression of activity dependent genes (ADGs).**

**(A)** t-SNE plot of subclustered MHb neurons extracted from cells in Figure 1B. **(B)** t-SNE plot showing three clusters of cells (top) that expressed elevated levels of several ADGs (*Fos, Fosb, Egr1, Junb, Dusp18*, etc.). **(C)** t-SNE plot after regressing out the principle component (PC) that included many of the ADGs shown in (B). Cells from clusters that were high in ADG expression were now intermingled with clusters that we defined by the spatial location of their DEGs (See also Figure 2C and D). **(D)** t-SNE plot showing ADG score following regressing out of the PC containing ADGs. **(E)** All 12 statistically significant PCs for the MHb neuron clusters shown above. PC number 4 (red) contained several ADGs. **(F)** The top 25 genes associated with PC4 (the ADG PC) contained several known ADGs highlighted in red.

**Figure S3 (related to Figure 2): Sample ISH images showing spatial distribution of selected differentially expressed genes in MHb. (A-J)** Sample ISH images from the Allen Institute Database showing selected differentially expressed genes for distinct transcriptionally defined neuronal subtypes in MHb. Gene name is in the upper right of each image and subregion where gene is enriched is on the left. Scale bar = 250μm.

**Figure S4 (related to Figure 2 and 3): Differentially expressed genes define distinct habenular subtypes. (A)** Left: Dendrogram for subclustering of all neurons shown in Figure 2 and 3. Right: Dot plot displaying expression of example differentially expressed genes used to identify each subtype of habenula neuron. The color of each dot (blue to red) indicates the relative expression of each gene whereas the dot size indicates the fraction of cells expressing the gene. Only representative genes are shown for entire list of DEGs see Table 3 and 5.

**Figure S5 (related to Figure 5): Cells from all 4 LHb subtypes project to both the VTA and DRN. (A)** Quantification of fluorescence coverage of single cells for FISH of selected enriched genes in LHb neurons that were positive for *RbV-N* following monosynaptic retrograde tracing from VGAT-IRES-Cre+ neurons in the VTA (left: n= 521 cells, 4 mice; center: n=742 cells, 4 mice; right: n= 167 cells, 2 mice). **(B)** Quantification of fluorescence coverage of single cells for FISH of selected enriched genes in LHb neurons that were positive for *RbV-N* following monosynaptic retrograde tracing from DAT-IRES-Cre+ neurons in the VTA (left: n= 233 cells, 3 mice; center: n= 233 cells, 3 mice; right: n= 103 cells, 3 mice). **(C)** Quantification of fluorescence coverage of single cells for FISH of selected enriched genes in LHb neurons that were positive for *RbV-N* following injection of RbV-GFP into the DRN (left: n= 163 cells, 3 mice; center: n= 133 cells, 3 mice; right: n=169 cells, 3 mice).

**Figure S6 (related to Figure 5): Quantification and genetic characterization of VTA starter cells from monosynaptic retrograde tracing. (A)** Left: Coronal section of injection site into VTA and starter cells location for Cre-dependent monosynaptic retrograde tracing experiments. Right: FISH for *RbV-N* to demonstrate the location of rabies infected cells in the VTA. **(B)** Left: Coronal section of injection site into DRN Cre-independent retrograde tracing experiments. Right: FISH for *RbV-N* to demonstrate the location of rabies infected cells in the DRN. **(C)** Left: Quantification of fluorescence coverage of single putative starter cells (*Cre*+ *and RbV-N+*) for FISH of *Cre* and *Slc17a6* in VTA neurons of the VGAT-IRES-Cre animals (n= 567 cells, 4 mice). Right: The proportion of putative starter cells that expressed *Slc17a6*. There is a subset of VGAT-IRES-Cre+ neurons in the VTA that co-express *Slc17a6* (Root et al., 2014). (D) Left: Quantification of fluorescence coverage of single putative starter cells (*Slc6a3*+ *and RbV-N+*) for FISH of *Slc6a3* and *Slc32a1* in VTA neurons of the DAT-IRES-Cre animals (n= 566 cells, 3 mice). Right: The proportion of putative starter cells that expressed *Slc32a1*. **(E)** Negative control for EnvA pseudotyping of the rabies virus (EnvA-RbV-GFP) showing a coronal section following injection of EnvA-RbV-GFP into the VTA without prior infection by AAV-TVA-mCh. Without co injecting AAV-TVA-mCh, EnvA-RbV-GFP cannot infect neurons, thus no GFP expression. Filled rectangles represent the mean and error bars are ±SEM.

**Figure S7 (related to Figure 2, 3, and 4): A map of habenula subregions based on single cell transcriptomic profiling.**

**(A)** Habenular subregions are outlined in black, MHb subregions are green and LHb subregions are magenta. The location of borders is a rough estimate of a boundary between transcriptionally defined neuronal subtypes and previous literature.

