## Supplementary figures and images for "Distinct neuronal subtypes of the lateral habenula differentially target ventral tegmental area dopamine neurons"

### supplemental figures

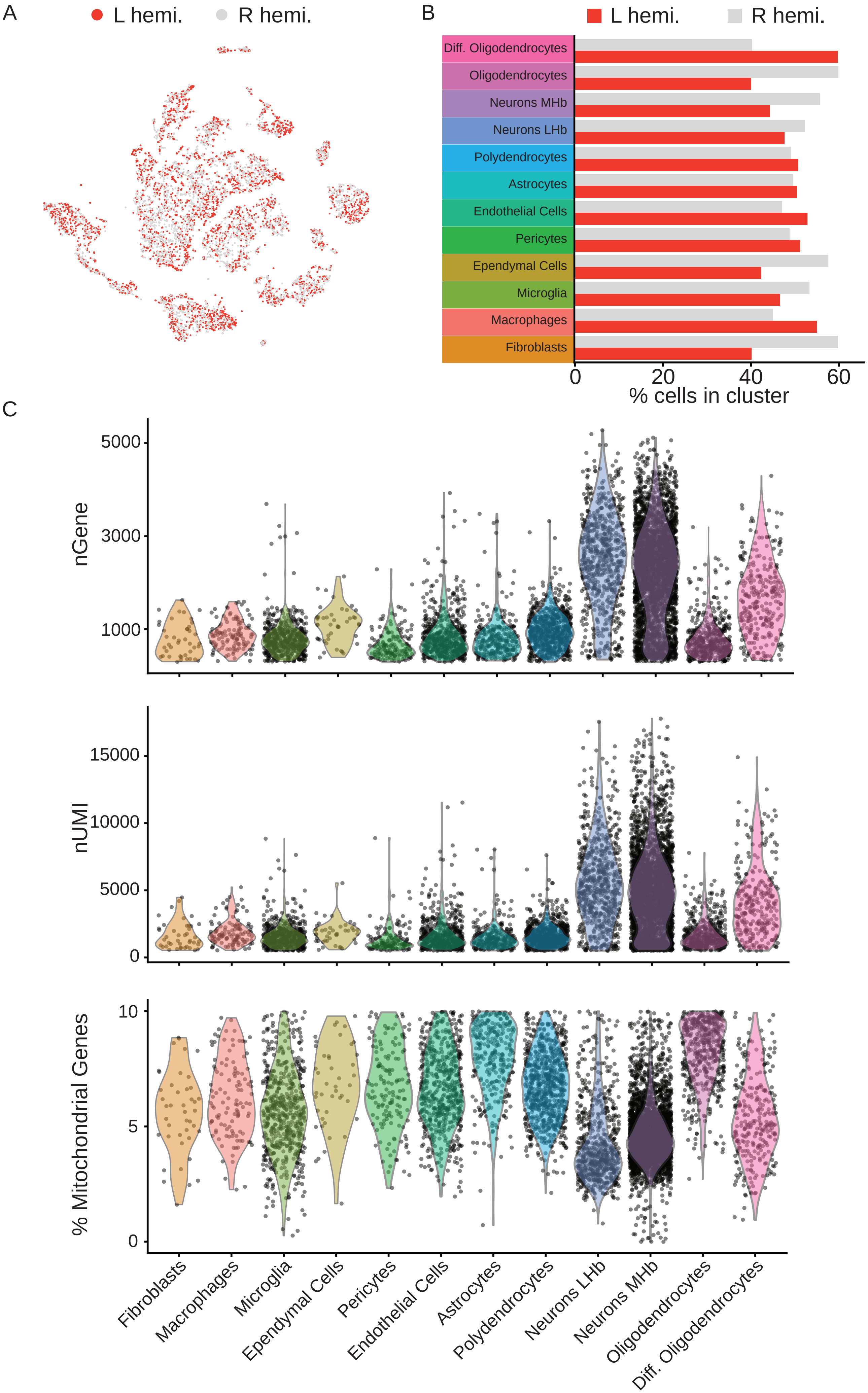

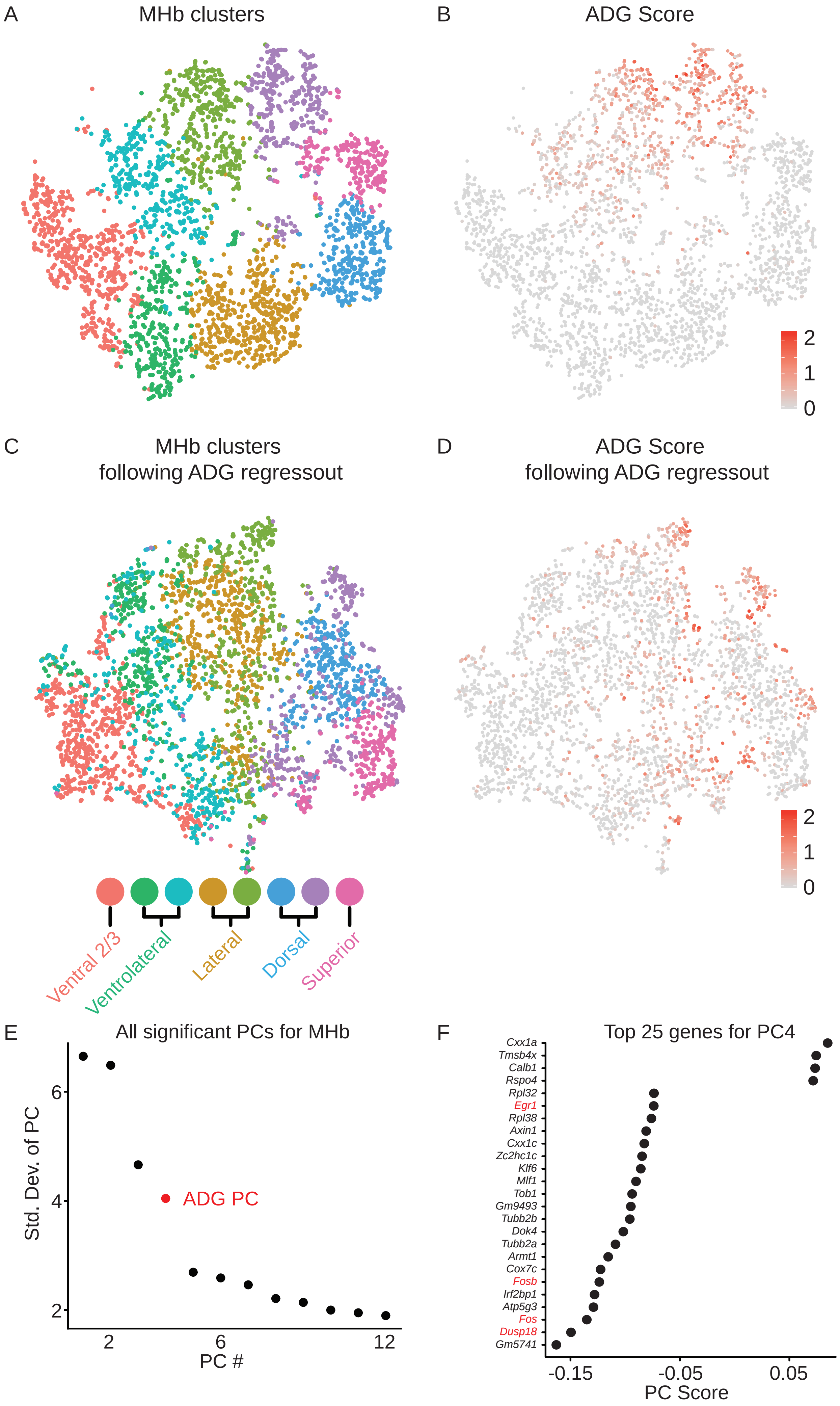

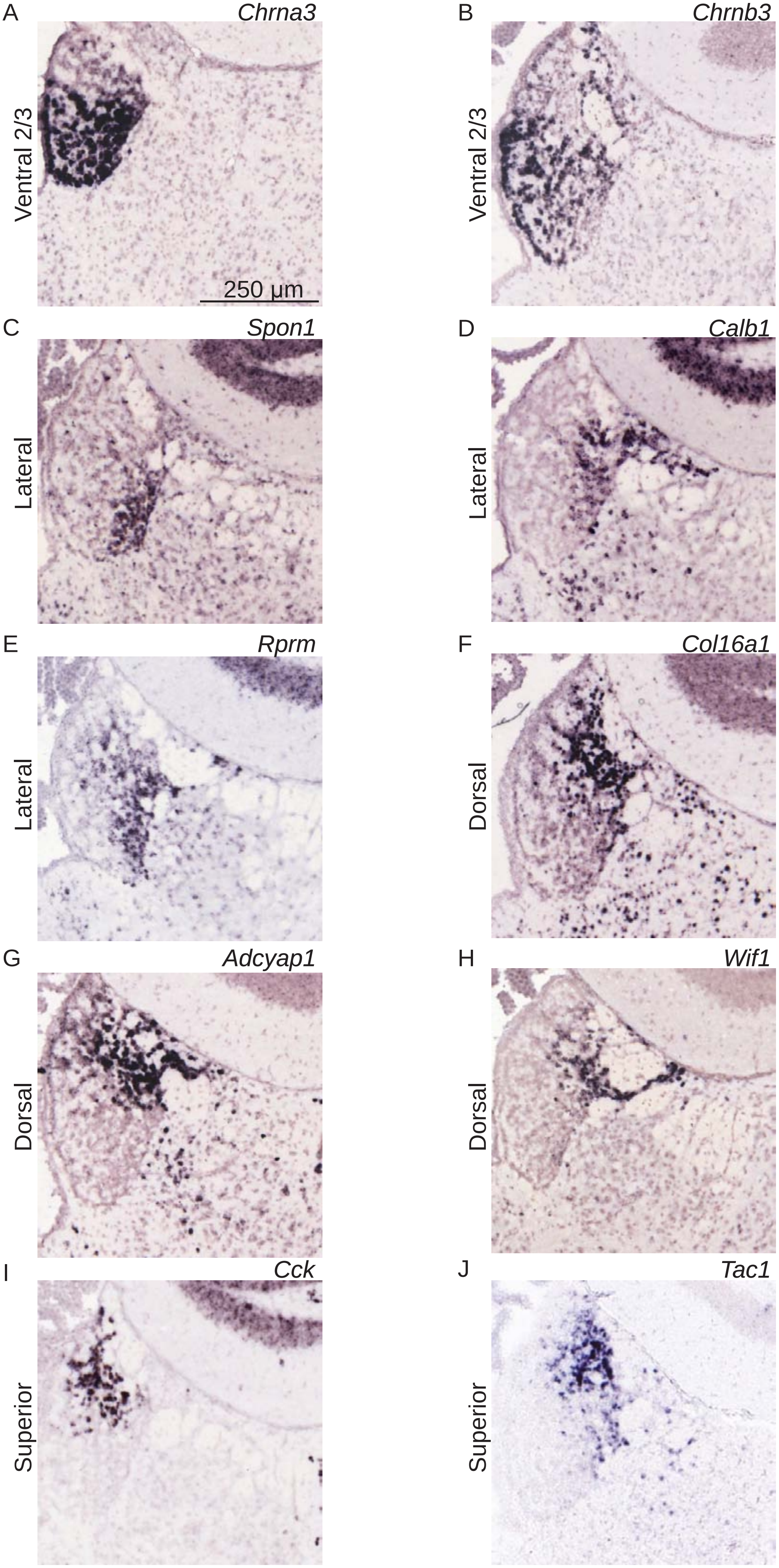

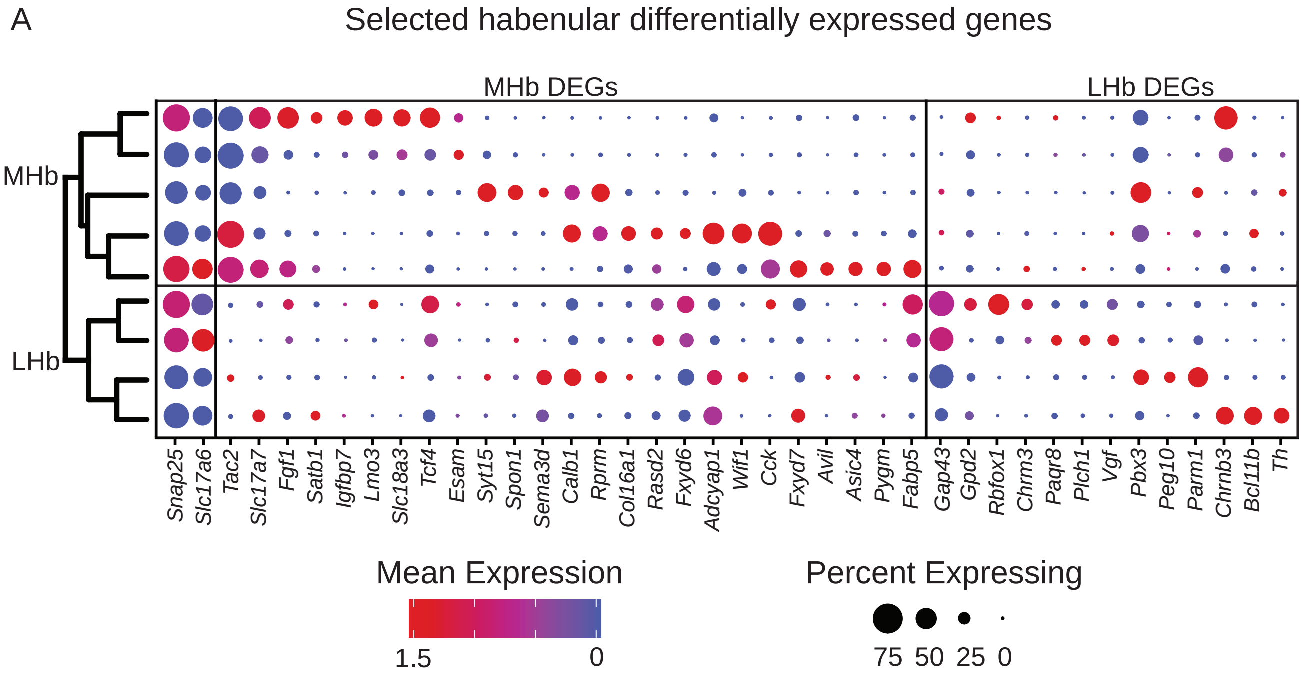

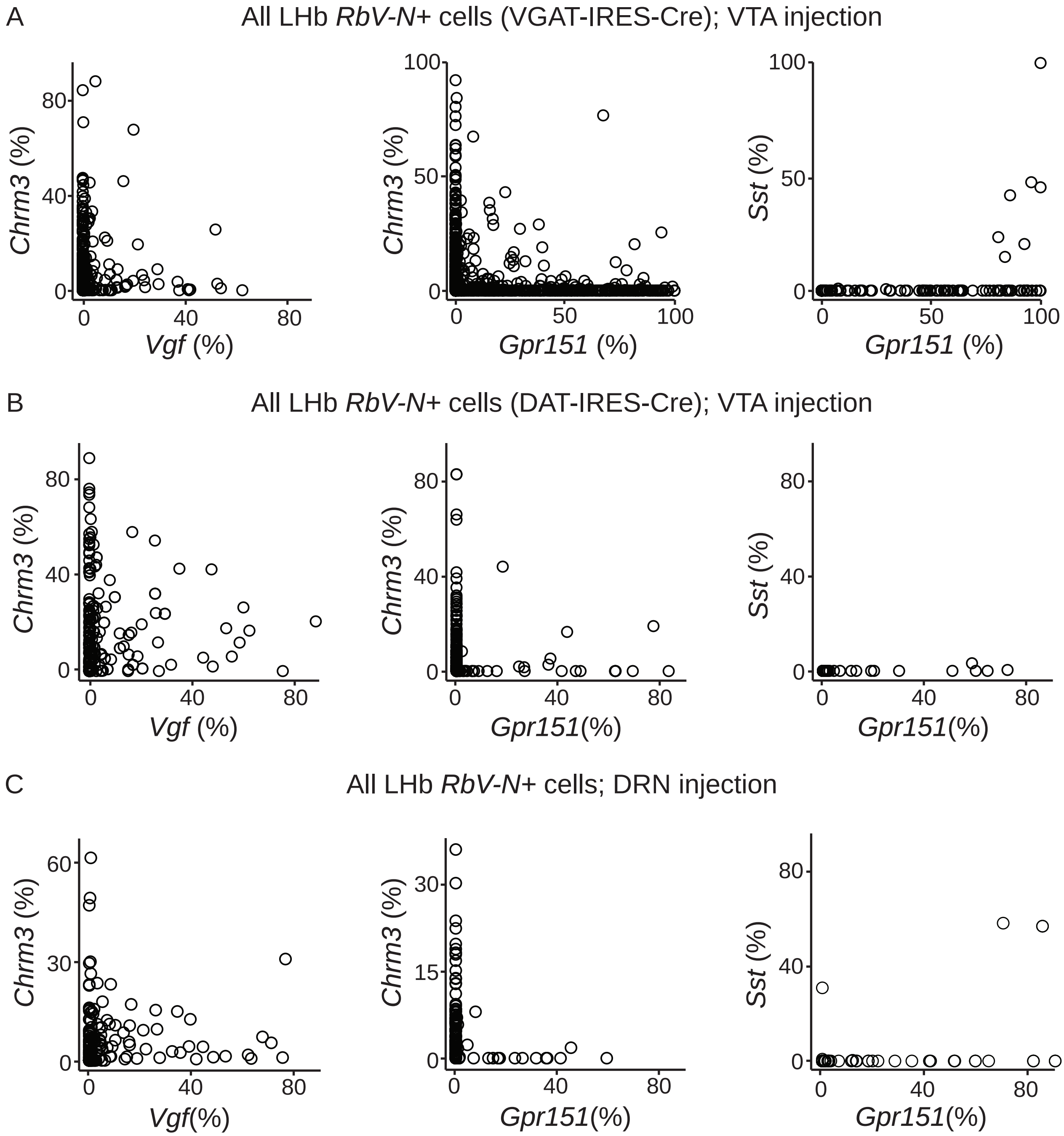

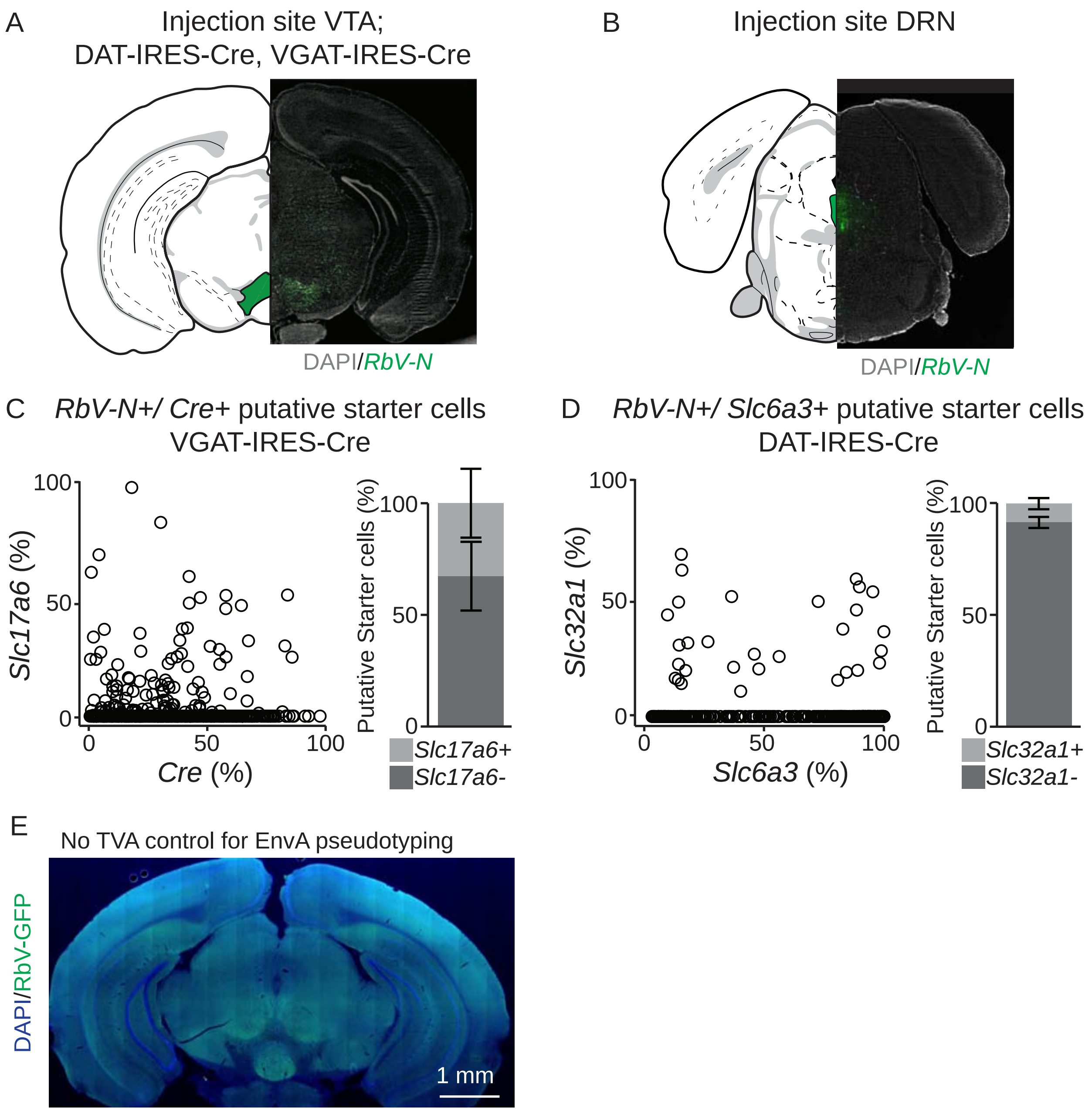

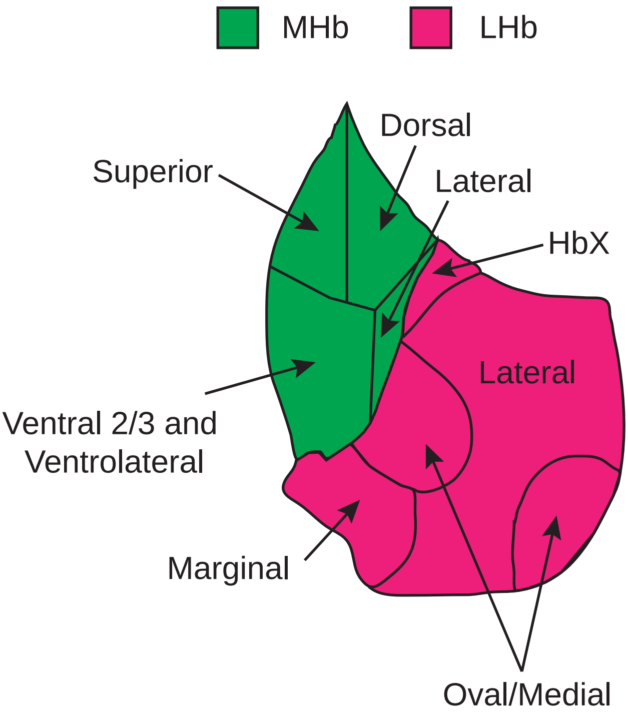
